# Transcranial focused ultrasound neuromodulation of the human primary motor cortex

**DOI:** 10.1101/234666

**Authors:** Wynn Legon, Priya Bansal, Roman Tyshynsky, Leo Ai, Jerel K. Mueller

**Affiliations:** Division of Physical Therapy and Division of Rehabilitation Science, Department of Rehabilitation Medicine, Medical School, University of Minnesota Minneapolis, MN, USA; Department of Neuroscience, University of Minnesota, Minneapolis, MN, USA; Department of Neurosurgery, School of Medicine, University of Virginia, VA, USA

**Keywords:** Ultrasound, primary motor cortex, transcranial magnetic stimulation, reaction time, human

## Abstract

Transcranial focused ultrasound is a form of non-invasive neuromodulation that uses acoustic energy to affect neuronal excitability. The effect of ultrasound on human motor cortical excitability is currently unknown. We apply ultrasound to the primary motor cortex in humans using a novel transcranial ultrasound and magnetic stimulation (TUMS) paradigm that allows for concurrent and concentric ultrasound stimulation with transcranial magnetic stimulation (TMS). This allows for non-invasive inspection of the effect of ultrasound on motor neuronal excitability using the motor evoked potential (MEP) generated by TMS. We test the effect of ultrasound on single pulse MEP recruitment curves and paired pulse protocols including short interval intracortical inhibition (SICI) and intracortical facilitation (ICF). We also test the longevity of the effect and the effect of ultrasound on the cortical silent period in a small sub-sample of participants. In addition, we test the effect of ultrasound to motor cortex on a stimulus response reaction time task. Results show ultrasound inhibits the amplitude of single-pulse MEPs and attenuates intracortical facilitation but does not affect intracortical inhibition. Early evidence suggests that ultrasound does not affect cortical silent period duration and that the duration of inhibition may be related to the duration of stimulation. Finally, ultrasound reduces reaction time on a simple stimulus response task. This is the first report of the effect of ultrasound on human motor cortical excitability and motor behavior and confirms previous results in the somatosensory cortex that ultrasound results in effective neuronal inhibition that confers a performance advantage.

---

Transcranial focused ultrasound (tFUS) is an innovative approach to noninvasive neuromodulation that uses focused mechanical energy to provide high spatial resolution targeting of distinct cortical areas (Legon et al. 2014; Tufail et al. 2010). Previous research has shown that tFUS modulates neuronal activity in mice (Tufail et al. 2010), rats (Younan et al. 2013), rabbits (Yoo et al. 2011), sheep (Lee, Lee et al. 2016), pigs (Dallapiazza et al. 2017) and monkeys (Deffieux et al. 2013). tFUS has also been demonstrated to be a safe and effective method for transient neuromodulation in human somatosensory (Lee et al. 2015; Legon et al. 2014) and visual cortex (Lee, Kim et al. 2016). In Legon et al. (2014) we demonstrated tFUS to inhibit the primary somatosensory cortex and for this neuromodulation to be spatially specific. In addition, we found ultrasound to improve tactile discrimination performance. Here, we extend these findings to the human primary motor cortex (M1). To assess the effect of ultrasound to M1 we have developed a novel method of transcranial ultrasound magnetic stimulation (TUMS) that allows for concurrent and concentric delivery of focused ultrasound with transcranial magnetic stimulation (TMS). This pairing is advantageous as it allows for assessment of ultrasound on established and well-understood TMS metrics like the motor evoked potential (MEP). In addition, it allows for a non-invasive examination of the effect of ultrasound on specific neuronal populations as different TMS methodologies (ex. paired-pulse) have been demonstrated to preferentially affect different motor microcircuits (Di Lazzaro et al. 2012). Here, we test the effect of focused ultrasound to M1 in five separate experiments. We test the effect of ultrasound on single pulse MEP recruitment curves, different paired-pulse inter-stimulus intervals (1-15 msec) as well as the cortical silent period. In addition, we test the duration of the effect by offsetting the timing of tFUS with single-pulse TMS and finally, test the effect of tFUS to M1 during a simple stimulus response reaction time task. Based upon previous literature assessing the effect of tFUS on small and large animal motor cortex that has found ultrasound to elicit peripheral motor responses (H. Kim, Chiu, Lee, Fischer, Yoo 2014; King et al. 2013; Lee et al. 2016; Tufail et al. 2010; Yoo et al. 2011; Younan et al. 2013), tFUS to human motor cortex could have excitatory effects and increase MEP amplitude; however, from our work in human somatosensory cortex we expect the overall effect to be for inhibition of motor cortical excitability. Using different TMS methods that probe different motor neuronal circuits in addition to assessing a simple motor task we hope to gain a better understanding of the effect of tFUS in human motor cortex and the neuronal circuits that ultrasonic mechanical energy affects.

## Materials and Methods

### Subjects

All experiments were conducted with the approval of the University of Minnesota institutional review board. A total of 57 healthy volunteers; 21 men and 36 women, aged 19 to 38 years (22 ± 3.59 years) participated in the five separate experiments. All subjects gave written informed consent and were financially compensated for participating. Subjects were screened for eligibility and confirmed to be physically and neurologically healthy with no history of neurological disorder. Additionally, subjects were cross-checked for known medication contraindications to other non-invasive neuromodulation techniques (Rossi et al. 2009).

## General Experimental Procedures for all experiments

General experimental procedures common to the five experiments are outlined below. Procedures specific to the individual experiments are elaborated in the individual experiment methods sections.

### Transcranial ultrasonic magnetic stimulation (TUMS)

The TUMS coil uses a standard commercial figure eight coil of two coplanar coil windings of equal size (double 70 mm alpha coil Magstim Inc., UK) to which a custom made low profile (1.25 cm height) single element 0.5 MHz focused ultrasound transducer (Y. Kim et al. 2014) is attached at the center of the coil intersection of the coil windings using a custom 3D printed holder (Fig. 1A). Initial testing was conducted to study the safety and feasibility of concurrent and concentric tFUS/TMS. Initial feasibility testing concentrated on the potential interaction of the two energy sources and potential damage or induction of current in the ultrasound transducer from the TMS magnetic field and for the ultrasound transducer to affect the magnetic field produced by the TMS coil and subsequent electric field produced in the head. To measure the effect of TMS on the resultant sound field we constructed a custom acoustic tank that separated the TMS coil from the water but left the transducer at center axis 1 mm from the face of the TMS coil. A hydrophone (HNR-0500 Onda Corp. Sunnyvale CA, USA) was placed in the tank at the measured focus of transducer. The ultrasound transducer used in the TUMS coil is a custom made single element focused 0.5 MHz transducer with a focal length of 22 mm, diameter of 36.5 mm and height of 12.5 mm. Testing consisted of 100 single pulses of 100% stimulator output TMS delivered at an inter-stimulus interval (ISI) of 8 seconds with and without concomitant ultrasound. The resultant TMS artifact and sound field were captured by the hydrophone in the test tank using test software and 3D stage (Precision Acoustics Dorset, UK). The TMS artifact observed by the hydrophone without the application of ultrasound was first recorded, followed by the pressure field from the US transducer (250 cycles) without TMS. Then, TMS was delivered concurrently with US at the same points to compare the pressures produced by the transducer with and without TMS. Traces of the TMS artifact alone were first subtracted from traces captured with simultaneous US and TMS and compared to the pressure from US alone. Next, we tested the effect of the US transducer on the magnetic field produced by the TMS coil. We measured the resultant magnetic vector potential from the center of the coil using a custom made magnetic probe made of two rectangular shaped windings of wires (1cm^2^ surface area) (Opitz et al. 2015) that were oriented perpendicularly to each other and placed in plane with the face of the ultrasound transducer 1.25 cm above the face of the TMS coil. A piece of paper outlining a 3x3 cm grid was placed in plane with the face of the US transducer above the TMS coil as a guide for measurement points of the magnetic vector potential. Single pulse TMS was delivered at 100% stimulator output at an ISI of 8 seconds and recorded in the two axes of the probe at all points of the grid both with and without the ultrasound transducer present at the center of the TMS coil. Comparison of the magnetic vector potential measurements at each of the points with and without the US transducer were conducted and compared between the transducer and no transducer using a paired t-test.

**Figure 1.**
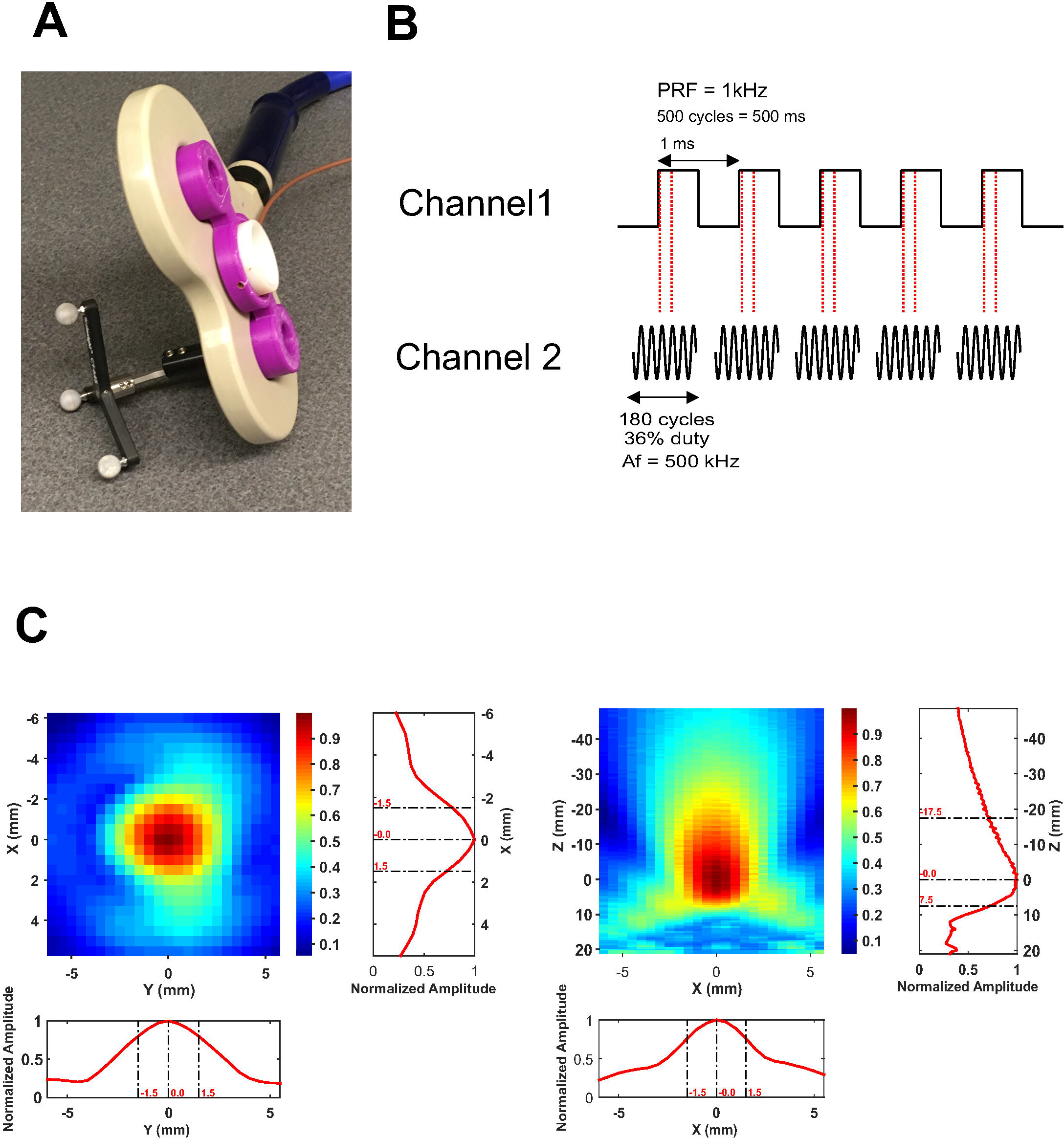
Transcranial ultrasound and magnetic stimulation (TUMS). **A.** Photograph of the TUMS device showing the TMS coil (beige), ultrasound transducer (white) and the holder (purple). Tracking bulbs are also visible and used to guide TUMS to a specific brain target using stereotaxic neuronavigation. **B.** Ultrasound pulsing strategy. PRF = pulse repetition frequency; Af = acoustic frequency. **C.** Pseudo-color free water plots of ultrasound pressure field. Scales are normalized to maximum pressure.

To test the impact of the standoff height of the transducer on the induced electric field in the head from TMS, we modeled the 70 mm figure eight coil this using SIMNIBS 2.0 (www.simnibs.de). We modeled two conditions: a monophasic posterior to anterior current with the 70 mm figure eight coil tangential to the scalp over the primary motor cortex with handle posterior angled at 45 degrees from midline resting against the scalp, and the other condition kept all parameters the same except had the TMS coil offset from the scalp by 1.25 cm; the height of the transducer.

### tFUS waveform and delivery

For all experiments we used a custom-made single element focused transducer with a center frequency of 0.5 MHz, 30 mm aperture and 22 mm focal length. The induced field in free water from this transducer is pictured in Fig 1C.

The waveform used was the same as previously described (Legon et al. 2014; J. K. Mueller, Ai, Bansal, Legon 2016)). This waveform was generated using a two-channel 2-MHz function generator (BK Precision Instruments, CA, USA). Channel 1 was set to deliver tFUS at a pulse repetition frequency (PRF) at 1 kHz and channel 2 was set to drive the transducer at 500 kHz in burst mode while using channel 1 as the trigger for channel 2. Channel 2 was set to deliver 180 cycles per pulse, and channel 1 was set to deliver 500 pulses, resulting in a 500 msec duration. Channel 2 output was sent to a 100W linear amplifier (2100L Electronics & Innovation Ltd, NY, USA), with the output of the amplifier sent to the custom made, low profile tFUS transducer (Fig 1B). For both the single pulse and paired pulsed experiments, ultrasound was time-locked to occur 100 msec prior to the TMS pulse. For all experiments, the ultrasound condition involved acoustically coupling the active face of the ultrasound transducer to the scalp at the pre-determined site (see stereotaxic neuronavigation). To achieve acoustic coupling to the head, the hair was parted to expose the scalp and ultrasound gel was used to keep the hair out of the way and ensure proper coupling with the ultrasound transducer. The transducer was also prepped with ultrasound gel on the surface that met the head, and was then placed on the exposed scalp as has been previously employed (Legon et al. 2014, Mueller et al. 2014, Ai et al. 2016). The sham condition involved turning over the transducer so that no ultrasound was delivered to the head, though contact was maintained ensuring equitable auditory artifact. This is important as in some cases, the piezoelectric element in the transducer produced a slightly audible buzzing. As such, participants were provided ear protection and did not report any sensible differences between sham and ultrasound for any of the experiments.

### Quantitative modeling of ultrasound wave propagation

A computational model was run to visualize and evaluate the wave propagation of tFUS across an example skull to determine the resultant intracranial beam profile and pressure map in the brain. The model was run using a magnetic resonance (MR) imaging and computerized tomography (CT) dataset taken from the Visible Human Project ® (Spitzer and Whitlock 1998). The transducer was placed on the scalp site overlying the hand knob of the primary motor cortex. Simulations were performed using the k-Wave MATLAB toolbox (Treeby and Cox 2010). Methods and details of the modeling parameters can be found in J. K. Mueller et al. (2017).

### Stereotaxic neuronavigation

Identification of all targeted areas for all experiments was aided and confirmed using a stereotaxic neuronavigation system (BrainSight, Rogue Research, Montreal, QUE, CAN) that was retrofitted to work with the TUMS coil and the individual tFUS transducer used in the reaction time experiment (see Figure 1A). Participants were fitted with infrared sensors to track head movement and individual fiducial head markers were used to align the participants’ head with a template brain in the neuronavigation system. The TUMS coil and the ultrasound transducer (reaction time experiment) were also fitted with infrared markers allowing for detailed position of the coil or transducer relative to the participant head. This aided in initial targeting the primary motor cortex but more importantly ensured constant placement of the TUMS coil and tFUS transducer trial to trial throughout the duration of all experiments. Target tolerance was set <= 1 mm from the original determined neuromodulation position.

### Electromyography (EMG)

For all experiments surface peripheral EMG was collected using surface adhesive electrodes (Medi-Trace ® 530 series) in a belly tendon montage ground to the medial epicondyle of the ipsilateral humerus. Electrodes were additionally secured in place with tape to ensure good continuous contact and no movement. For experiments 1-4 we recorded from the first dorsal interosseous (FDI) of the dominant hand. For experiment 5 (reaction time) we recorded from the dominant abductor pollis brevis (APB) as this was the muscle performing the task. EMG data was continuously recorded using a DC amplifier (Net Amps 400, Electrical Geodesics, Inc. Eugene, OR, USA) sampled at 1kHz. Data was stored on a PC for later offline analysis. EMG analysis was performed using custom-made scripts written in Matlab (Mathworks Natick MA USA). The continuous data was first filtered forwards and backwards to avoid edge artifacts and to eliminate phase shifting using a 3^rd^ order Butterworth bandpass filter with cutoffs 5 – 200 Hz. The data was then epoched −50 to 100 msec around the onset of TMS delivery. Motor evoked potential (MEP) amplitudes were identified visually and quantified using the peak to peak method. An MEP was considered to be present if easily identifiable peaks were apparent above the noise floor of that trial in a 50 millisecond time window post TMS onset. Events where no MEP was identified were given an amplitude of zero and included in subsequent data analysis.

### Targeting of M1 hotspots

Targeting of M1 was similar for each of the five experiments. For each participant, identification of the MEP ‘hotspot’ was determined by first setting the TMS stimulator output to 50% of stimulator maximum. The TMS coil (no ultrasound transducer) was positioned over the contralateral somatomotor region with the handle positioned backwards at 45 degrees from midline for a posterior to anterior induced current. Single pulses in sets of five (ISI = 10 seconds) were delivered and stimulator intensity adjusted between sets until MEPs at least 50 µV were elicited for three of the five stimulations. Subsequent target areas were tested spaced 1 cm anterior, posterior, lateral and medial to the initial target to verify the lowest possible stimulator output to meet threshold criteria. If one of these targets proved to have a lower threshold than the initially identified target, a new set of five single pulses was delivered to areas 1 cm anterior, posterior, lateral and medial to this new spot to ensure the lowest threshold of stimulation. This process was repeated until the site of lowest intensity was determined. Spacing of 1 cm was used as this was double the lateral spatial resolution of the ultrasound transducer used and thus provided spatially discrete (non-overlapping) sites for potential hotspot location. An additional step was necessary as the TUMS device produces a different resting motor threshold (RMT) as compared to the TMS coil alone (due to the standoff height of the ultrasound transducer). At the pre-determined hotspot as determined above, RMT for TUMS was determined using sets of ten stimuli (ISI = 10 seconds) and the stimulator output was set at a percent stimulator output that resulted in MEPs of at least 50 uV 50% of the time. In general, the TUMS RMT was about 20-30% higher than the RMT of the TMS coil alone. In cases where the TUMS RMT resulted in a stimulator output > 100%, further study was not possible and the participant did not continue in the study.

### Experiment 1: Single-Pulse MEP recruitment curves

#### Subjects

A total of twelve individuals (4 male, 8 female, 22 ± 3.59 years) participated in this experiment. All were self-reported right hand dominant.

#### Experimental procedures

To generate recruitment curves, individual resting motor threshold for the dominant first dorsal interosseous (FDI) was determined as above for the TUMS coil and a stimulator output 20% below this value (rounded to the nearest 5%) was used as the starting value. For example, if a participant had a RMT of 73% stimulator output, their initial testing point would be 55% stimulator output (73-20 rounded to nearest 5%). Testing increments were conducted beginning at the above calculated starting stimulator intensity and increased in increments of 5% until 100% stimulator output was reached. A total of ten single pulses were collected for each stimulator output at an ISI of 10 seconds. Prior to collection of the recruitment curve, a baseline condition (no ultrasound, but with 1.25 cm standoff) was collected of 10 single pulses spaced 10 seconds apart at individual TUMS RMT. This procedure was completed for both tFUS and sham conditions, counterbalanced across participants. Stimulation time was approximately 15-20 minutes for each condition, and total experiment time including setup was approximately 2 hours.

#### Analysis

To test for significant differences in the MEP recruitment curves between sham and TUMS neuromodulation an analysis of covariance (ANCOVA) was conducted with stimulation intensity as the covariate (Wassermann et al. 1998). The independent variable was the condition (sham vs. TUMS). For this analysis, only data points from stimulator intensities 75 – 100% were included as these were the stimulator intensities that generally produced non-zero quantifiable MEPs across participants that could be appropriately modeled using linear regression.

### Experiment 2: Paired-Pulse MEP

#### Subjects

A total of ten individuals (3 male, 7 female, 22 ± 3.24 years) participated in this experiment. All were self-reported right hand dominant.

#### Experimental procedures

The paired pulse technique for this study was adapted from (Kujirai et al. 1993) where a sub-threshold conditioning stimulus is followed by a supra-threshold test stimulus delivered through the same coil at a given latency. Prior to paired-pulse testing, TUMS RMT was established so as to determine the 80% and 120% TMS stimulator outputs for paired pulse testing. A baseline condition was then completed as ten single pulses at 120% TUMS RMT spaced at 10 seconds. After baseline collection, the TUMS device was positioned over the dominant M1 FDI hotspot with the handle pointing posterior at 45 degrees to midline. The conditioning stimulator was set to 80% TUMS RMT while the test stimulator was set to 120% TUMS RMT. Ten TUMS stimulations were delivered at an ISI of 10 seconds for each inter-stimulus interval (ISI) of 1-15 msec. The order of the paired pulse ISIs was always ascending from 1 to 15 msec, and this procedure was completed for both tFUS and sham conditions, the order of which was counterbalanced across participants. Stimulation time was approximately 25-30 minutes for each condition, and total experiment time including setup was roughly 2.5 hours.

#### Analysis

Data were first normalized to individual TUMS 120% RMT baseline. The data set was parsed into data windows that represent established ISIs for short intracortical inhibition (SICI) (1-5 msec) and intracortical facilitation (ICF) (10-15 msec) (Kujirai et al. 1993) to test for specific effects of ultrasound on these mechanisms. The intermediate data section (6-9 ISI) was also analyzed separately though these ISIs have not been demonstrated to be specific for either inhibition or facilitation using the paired pulse paradigm. For each data epoch, a separate two-way repeated measures analysis of variance (ANOVA) was conducted with factors MODULATION (sham, ultrasound) and TIME (ISI times). Post-hoc Tukey tests were conducted to investigate any significant effects.

### Experiment 3: Cortical Silent Period (CSP)

#### Subjects

A total of four (3 male, 1 female, 28.4 ± 5.9 years) participated in this experiment.

#### Experimental procedures

To preliminarily test the effect of tFUS on the CSP we delivered TUMS during active muscle contraction of the dominant FDI muscle. Participants were required to pinch a dynamometer (HD-BTA, Vernier, OR, USA) fixed to a table top positioned such that a comfortable thumb index finder pinch could be performed to achieve activation of the FDI muscle. FDI activation was confirmed prior to formal testing. Individual participant maximum FDI force for this pinch task was collected prior to formal testing and participants were required to maintain 30% of this value during experimentation (Werhahn et al. 1999). To ensure this, real-time biofeedback of their FDI force output was provided on a computer screen positioned in front of them using custom scripts written in LabView (National Instruments, TX, USA) that used a dial and needle to indicate the force. A tolerance of ± 3% of the target force was used. If participants force was outside this range, a red light indicated they should adjust their force to remain in the acceptable range. Prior to formal testing, a baseline condition of 10 TMS single pulses spaced 10 seconds apart with no ultrasound was collected using the TUMS device positioned with the handle pointing backwards at 45 degrees to midline over the FDI hotspot at 120% RMT while subjects held the tonic contraction. During testing, the TUMS device was positioned similarly over the FDI hotspot and set to 120% of resting motor threshold. Ten stimulations were delivered for both tFUS and sham counterbalanced across participants at an ISI of 10 seconds while participants continuously held the 30% contraction.

#### Analysis

Quantification of silent period duration was done visually using custom made scripts written in Matlab (Mathworks, Natick, MA, USA). Data was first filtered using a forward and reverse Butterworth 3rd order bandpass filter with 5 – 200 Hz cutoff and then rectified. Data were epoched −50 to 100 milliseconds around the TMS event. We used visual MEP onset to EMG return for measurement as recommended by (Damron et al. 2008). Durations were averaged (n = 10) for each condition for each participant and subjected to a one-way repeated measures analysis of variance (ANOVA) to test for statistical differences between conditions (baseline, sham and TUMS). In addition to CSP duration, we also quantified MEP amplitude during contraction and tested for condition differences using a one-way repeated measures ANOVA.

### Experiment 4: Duration

#### Subjects

A total of four (3 male, 1 female, 28.0 ± 6.6 years) participated in this experiment.

#### Experimental Procedures

To preliminarily test the longevity of the effect of tFUS on motor cortical excitability we delivered single pulse TMS at different time points relative to the offset of 500 milliseconds of tFUS. It was the purpose of this experiment to better understand the lasting effects of ultrasound on motor cortical excitability. Our previous work reported no discernable cumulative effects of repeated ultrasound neuromodulation on potentials of the somatosensory evoked potential (Legon et al. 2014) and here, using TUMS, we can better understand if ultrasound induced neuromodulatory effects persist beyond the duration of the ultrasound stimulation on a trial to trial basis. To do so, we delivered ultrasound at varying time offsets preceding single pulse TMS. The time offsets included −100 msec, 100, 200, 500, 1000, 2000, 5000 and 10000 milliseconds relative to the TMS single pulse onset. Prior to collection of each time offset, a baseline condition was also collected where no ultrasound was delivered during single pulse TMS. For each offset, a total of 10 single pulses (ISI = 10 seconds) were collected for the TUMS condition and the order of the offsets was randomized across participants.

#### Analysis

As in previous experiments, EMG data was first filtered, epoched and MEP amplitude was quantified visually using the peak to peak method as previously described. Epochs that did not contain a discernable MEP were given a zero amplitude and included in subsequent analysis. Prior to statistical analysis, data from each time epoch was normalized to individual baseline values. Each time epoch was compared to the baseline condition using a paired t-test.

### Experiment 5: Reaction Time

#### Subjects

A total of 28 participants (9 male, 19 female) with a mean age of 22 ± 1.71 years were included in this experiment. Three of the subjects were self-reported left hand dominant.

#### Experimental Procedures

Based upon the above physiological results, we subsequently tested if there is any behavioral result of ultrasound delivered to the primary motor cortex. Specifically, we tested the effect of tFUS on reaction time using a simple stimulus-response task. Participants were required to attend to visual stimulus on a computer screen that consisted of large white block X or O presented on a black background in the middle of the screen. Participants were required to press the space bar on a standard computer keyboard with their dominant thumb when they saw an X and to withhold a response when an O was presented. A total of 100 stimuli were presented at random time intervals between 3 – 6 seconds with 20% O trials. Ultrasound was time-locked to occur 100 msec prior to the visual stimulus. Participants responded with their thumb while recording EMG activity from the abductor pollicis brevis (APB) which produced a reliable EMG signal during the task. The APB hotspot was determined for every participant using TMS as previously described. We did not use the TUMS coil for this experiment but did employ the same ultrasound transducer with the same parameters as used in the above experiments. The transducer was positioned over the contralateral APB hotspot and held in place with a secure headband. Three conditions were collected. Real and sham neuromodulation were delivered to the APB hotspot and an active control condition was also collected where tFUS was delivered with identical parameters and timing to the vertex of the head as determined using international 10-20 EEG electrode placement site CZ. The order of conditions was randomized across participants.

#### Analysis

Reaction time was defined as the difference between the timing of the onset of the visual stimulus and the timing of the key press as recorded by custom made scripts written in Matlab (Mathworks, Natick, MA, USA). Trials where no key press was recorded or fell outside 1000 msec from stimulus onset or a wrong key was pressed were not included in subsequent analysis. Only trials where the stimulus was an X were included in the reaction time analysis. Trials where the stimulus was an O were included in the catch trial analysis. A correct catch trial response was no key press within 1000 msec of the visual stimulus onset. Any key press within this time window was considered an error and included in the analysis. The average reaction time was calculated from the acceptable trials for each condition (active sham, M1 sham, tFUS) for each participant and this was tested for statistical significance using a one-way repeated measures analysis of variance (ANOVA). For catch trial data, the number of correct withholding of a key press was expressed as a percentage of total (20) opportunities for each condition for each participant. This data was subjected to a one-way repeated measure analysis of variance (ANOVA) to test for statistical significance between the conditions.

## Results

### TUMS coil

To ensure no interaction of the energy fields during TUMS we measured the resultant ultrasound pressure profile with and without a concomitant TMS pulse and measured the resultant magnetic vector potential from the TMS coil with and without the ultrasound transducer attached to the face of the TMS coil. We found negligible differences in the amplitude or morphology of the ultrasound waveform with and without a TMS pulse repeated over 100 trials at a TMS stimulator output of 100%. Figure 2A shows a representative trace of the sound field with and without TMS. For 100 TMS pulses with and without ultrasound we found no differences in the magnetic field produced by the TMS coil t(8) = 0.45, p = 0.91 (see Figure 2B). We repeatedly tested the device at weekly intervals and found no effect on the resultant sound field or the magnetic vector potential with repeated use. We also modelled the resultant electric field in the brain for the TMS coil and the TUMS coil (1.25 cm height offset) to assess the effect of the standoff height of the ultrasound transducer. Increasing the standoff height of the coil 1.25 cm from the head weakens the strength of the induced electric field to roughly 70% (4.71 vs. 3.31 V/cm^2^) of the field produced with the TMS coil directly on the scalp and increases the spread of the induced electric field at the level of the cortical surface holding constant the current fed through the TMS coils (Figure 3A). The peak induced electrical field in the cortex is still located directly below the center point below the two windings. Thus, by compensating for the loss in electric field strength by increasing the current fed through the coils, the electric field strength at the level of the cortical surface can be maintained above threshold to produce a peripheral muscle response. This is indicated in Figure 4 where the average % TMS output to produce a MEP amplitude of ~ 50 µV was ~75% on average.

**Figure 2.**
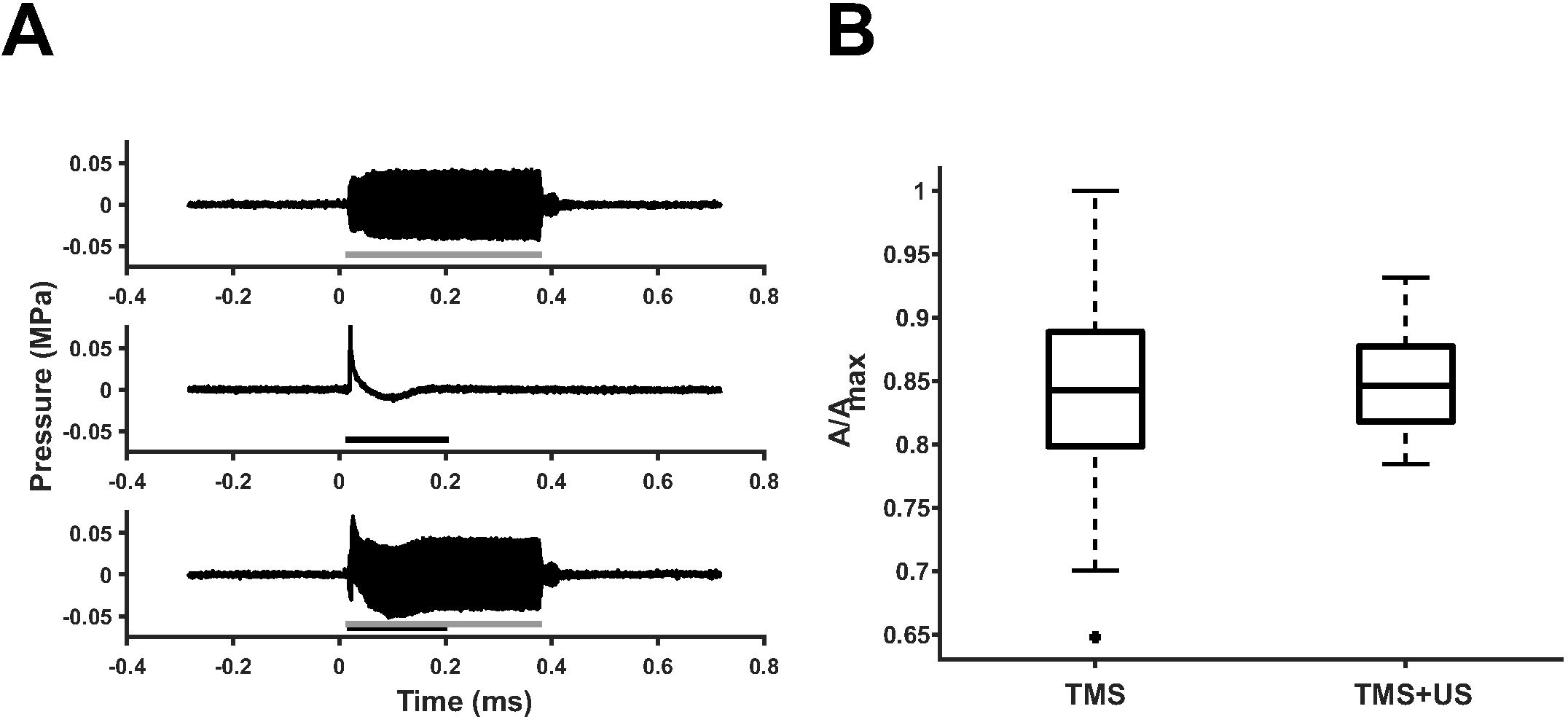
Interaction of ultrasound and TMS. **A.** Ultrasound field as measured in free water. Grey line represents on time of ultrasound. (Middle) Artifact from single 100% TMS stimulator output pulse. Black line depicts length of pulse artifact. (Bottom) Both ultrasound and TMS are delivered at the same time to determine impact of TMS on the sound field pressure. No effect of the TMS pulse was found on the sound field. The US transducer does not impact the amplitude of the TMS pulse. msec = milliseconds; MPa = megapascals. **B.** Average magnetic vector potential (A) relative to maximum (Amax) as measured from the plane of the face of the TMS coil without the ultrasound transducer (TMS) and with the ultrasound transducer affixed to the TMS coil (TMS + US). There was no difference in the average magnetic vector potential with the addition of the ultrasound transducer. The central line is the median, the edges of the box represent the 25% and 75% percentiles and the whiskers extend to the most extreme data points.

### Acoustic modelling

Figure 3B shows an example resultant modelled acoustic field using CT and MRI data directed at the hand knob of the primary motor cortex. These models take into consideration tissue properties that can affect the propagation of ultrasound for neuromodulation (J. K. Mueller, Ai, Bansal, Legon 2016; J. K. Mueller et al. 2017). It is evident from Figure 3B that there is good maintenance of the geometry of the sound field and that it is localized to the pre-central gyrus.

**Figure 3.**
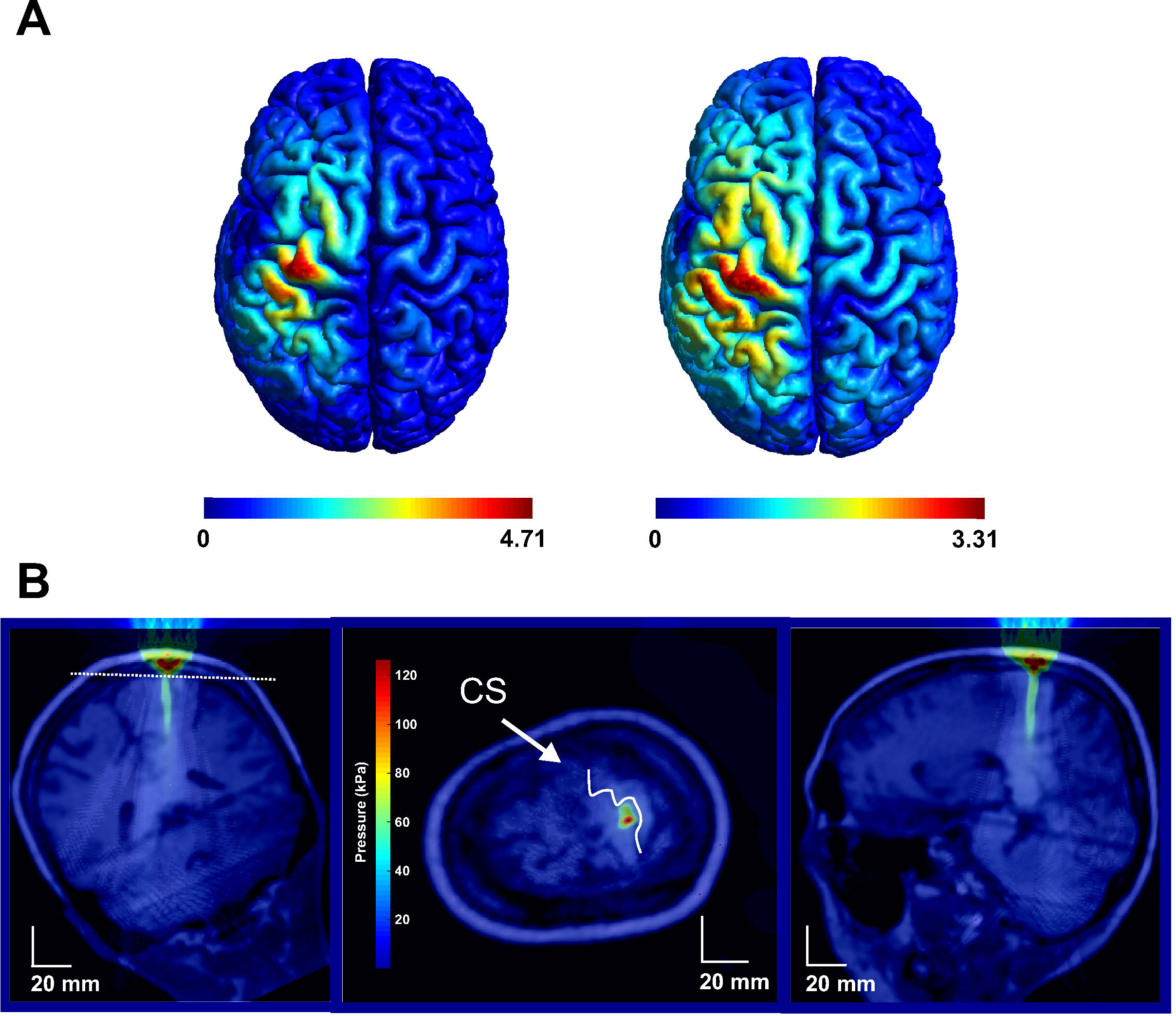
Modelled TUMS field and ultrasound field in the brain. **A.** Modelled induced electric field in the human brain from the TMS coil without the ultrasound transducer (left) and with the ultrasound transducer (right). **B.** Acoustic model of the ultrasound waveform positioned over primary motor cortex. Note: Middle figure is transverse section taken through the plane (white dashed line) in the left coronal figure. kPa = kilopascals.

### Experiment 1: Single pulse recruitment curves

The analysis of covariance revealed the slopes of the regression lines to be significantly different (F(1,8) = 5.28, p = 0.0406) between TUMS and sham indicating that the recruitment curves are different (Figure 4A). TUMS resulted in a general inhibition of MEP amplitude as stimulator percent increased as compared to sham stimulation. See Figure 4B for the raw recruitment data.

**Figure 4.**
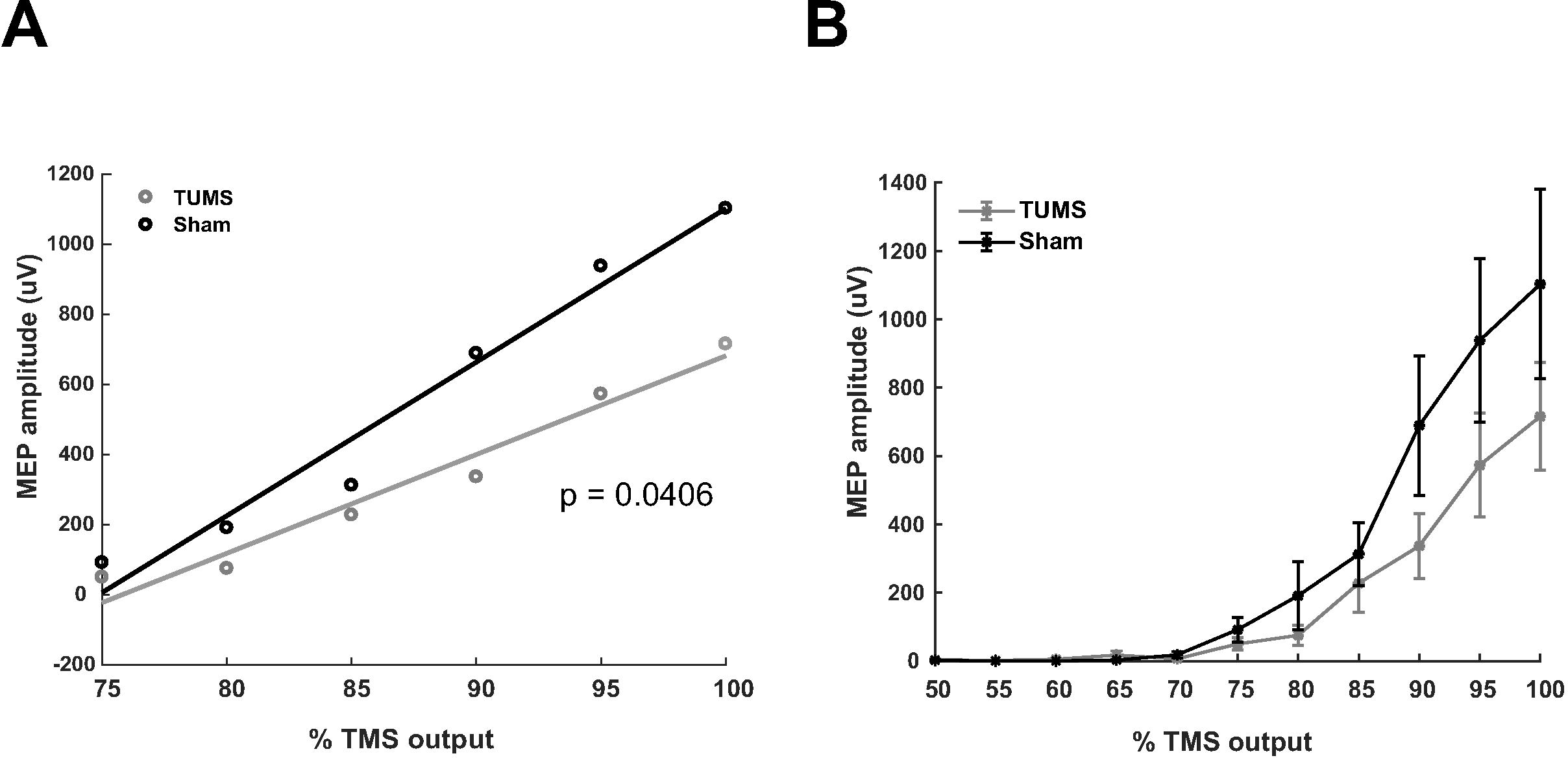
Ultrasound effects on single pulse TMS recruitment curves. **A.** Fitted linear regression curves for TUMS (grey) and sham (black) neuromodulation. The curves were statistically different (p = 0.0406). **B.** Group average (N = 12) raw data ± SEM recruitment curves for TUMS (grey) and sham (black) neuromodulation.

### Experiment 2: Paired-pulse

Baseline single pulse 120% TUMS RMT was 340.17 ± 47.27 µV. The two-way repeated measures ANOVA for SICI did not reveal a main effect for TIME F(4,90) = 2.15, p = 0.08 or MODULATION F(1,90) = 2.03, p = 0.1574. There was also no significant interaction F(4,90) = 0.29, p = 0.88 (Figure 5A & B). For the middle epoch consisting of ISIs 6-9 milliseconds the two-way repeated measures ANOVA did not reveal statistically significant effects for TIME F(3,72) = 0.62, p = 0.60, MODULATION F(1,72) = 1.11, p = 0.29 or the interaction F(3,72) = 0.23, p = 0.88 (Figure 5A & B). For the ICF epoch there was a significant main effect of MODULATION F(1,108) = 7.56, p = 0.007, no effect of TIME F(5,108) = 0.21, p = 0.95 and no interaction F(5,108) = 0.03, p = 0.99 (Figure 5A & B). The post-hoc Tukey test of the main effect of MODULATION revealed a significant difference for all ISIs 10 – 15 (p < 0.05) attenuated as compared to sham stimulation (Figure 5A).

**Figure 5.**
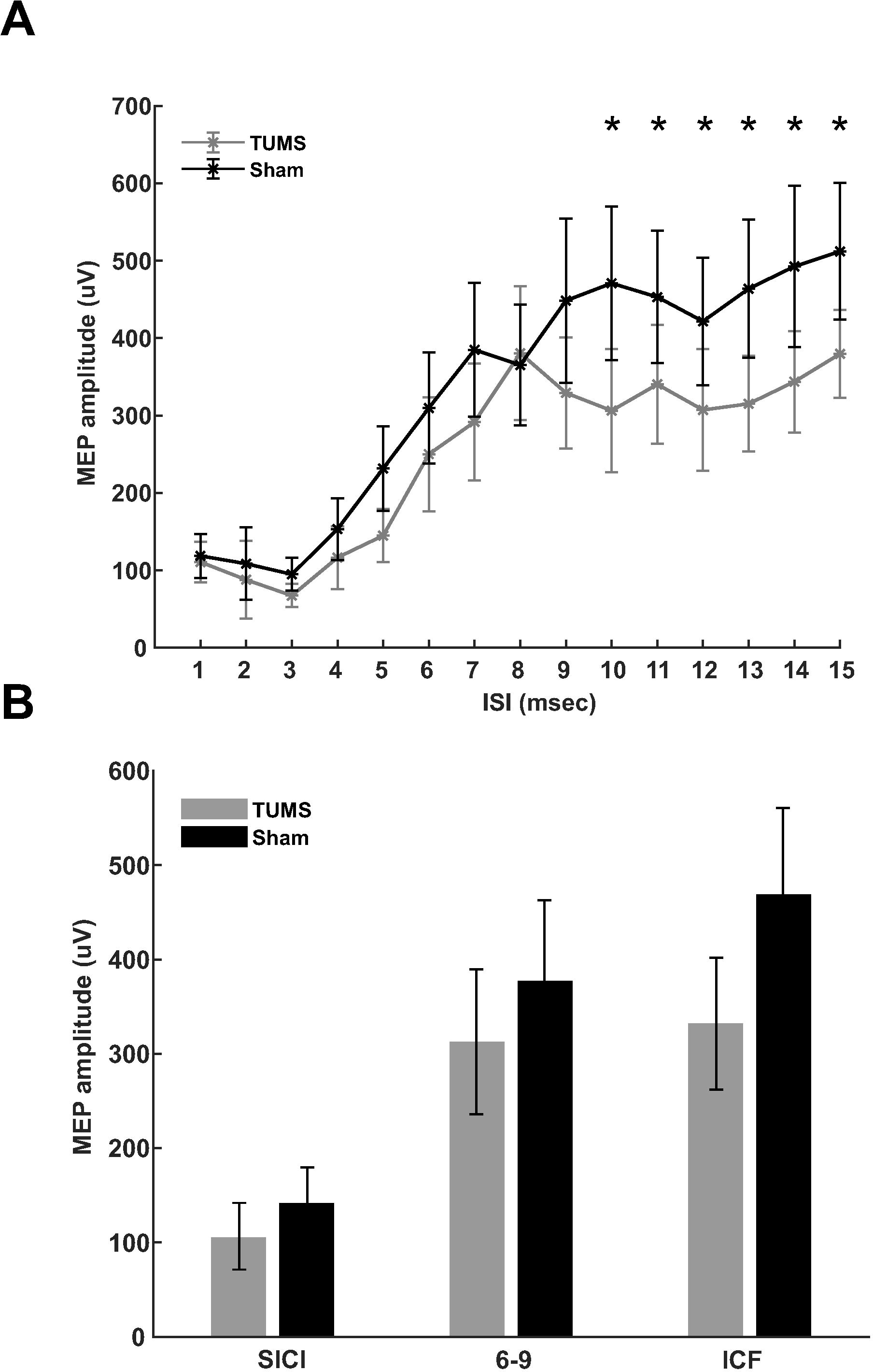
Ultrasound effects on paired-pulse TMS. **A.** Raw group average (N = 10) paired pulse average motor evoked potential (MEP) amplitude ± SEM for inter-stimulus intervals (ISI) 1 – 15 milliseconds. Transcranial ultrasonic and magnetic stimulation (TUMS) (grey) and sham (black) neuromodulation. Amplitudes are presented relative to a single pulse baseline (horizontal dashed line). * represents p < 0.05. **B.** Averaged normalized MEP amplitude ± SEM collapsed across grouped ISIs known to reflect short intracortical inhibition (SICI) and intracortical facilitation (ICF). * represents p < 0.05.

### Experiment 3: Cortical silent period

CSP durations were 71.12 ± 0.77 msec, 72.8 ± 1.2 msec and 70.66 ± 1.11 msec for the baseline, TUMS and sham conditions respectively. The one-way repeated measures ANOVA revealed no effect of TUMS on the duration of the cortical silent period (F(2,9) = 0.11, p = 0.89). The one-way repeated measures ANOVA revealed no effect of TUMS on the amplitude of the MEP during contraction (F(2,9) = 0.01, p = 0.99).

### Experiment 4: Duration of effect

We performed preliminary testing of the duration of the effect of ultrasound on motor cortical excitability as assessed by the peak to peak amplitude of the MEP at time points (-100, 100, 200, 500, 1000, 2000, 5000 and 10000 msec post ultrasound application. The average MEP peak to peak amplitudes of the MEPs were 0.76% ± 0.09%, 0.84% ± 0.13%, 0.85% ± 0.1%, 0.88% ± 0.12%, 0.98% ± 0.17%, 1.06% ± 0.17%, 0.99% ± 0.19% and 0.97% ± 0.08% for each time point respectively. The only time point to reach statistical significance was the −100 msec (t(3) = 3.23, p = 0.0484) (Figure 6).

**Figure 6.**
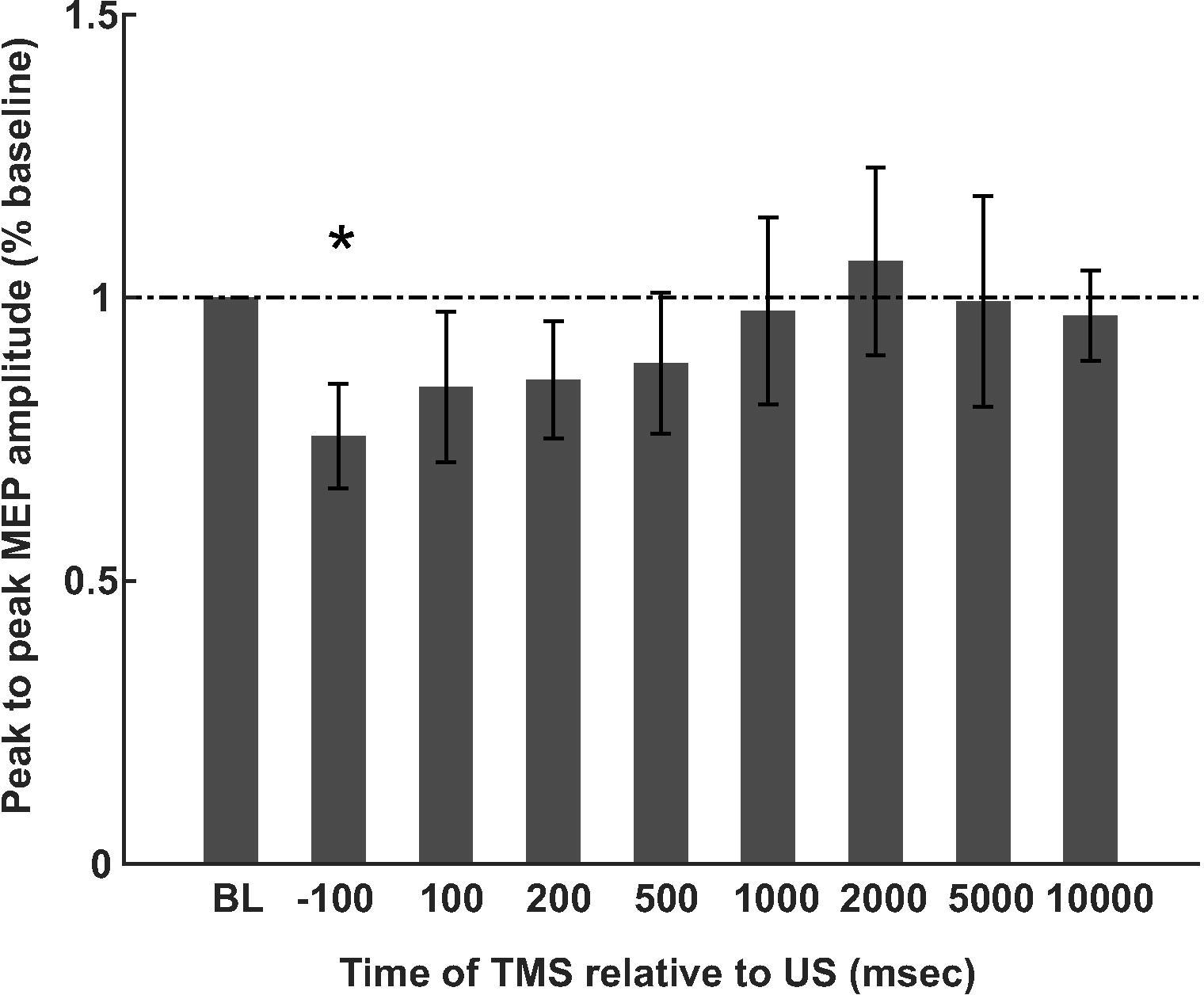
Longevity of ultrasound neuromodulation. Group average (N = 4) analysis of the duration of the effect of 500 milliseconds of pulsed ultrasound on motor evoked potential (MEP) amplitude expressed as a percentage of baseline. MEP excitability was tested out to 10000 milliseconds after the offset of ultrasound. * denotes p < 0.05.

### Experiment 5: Reaction Time

It was the purpose of this experiment to test if tFUS to the APB muscle representation during performance of a stimulus response reaction time task performed with the APB affected reaction time. Mean reaction times were 390.1 ± 54.2 msec, 389 ± 61.1 msec and 375.7 ± 58.6 msec for conditions active sham, sham and tFUS respectively. The one-way repeated measures ANOVA revealed a significant effect of condition F(2,54) = 3.96, p = 0.0248. Post-hoc Tukey test revealed the tFUS condition reaction time to be significantly lower than both the active and passive sham (p<0.05) (Figure7A). In addition to assessing reaction time, we were also able to assess performance on the task using the catch trial data. The percent correct responses to the catch trials were 91.9% ± 8.3%, 90.7% ± 8.5% and 92.5% ± 8.7% for the active sham, sham and tFUS conditions respectively. The one-way repeated measures ANOVA revealed no significant difference between conditions; F(2,54) = 1.04, p = 0.36 (Figure 7B).

**Figure 7.**
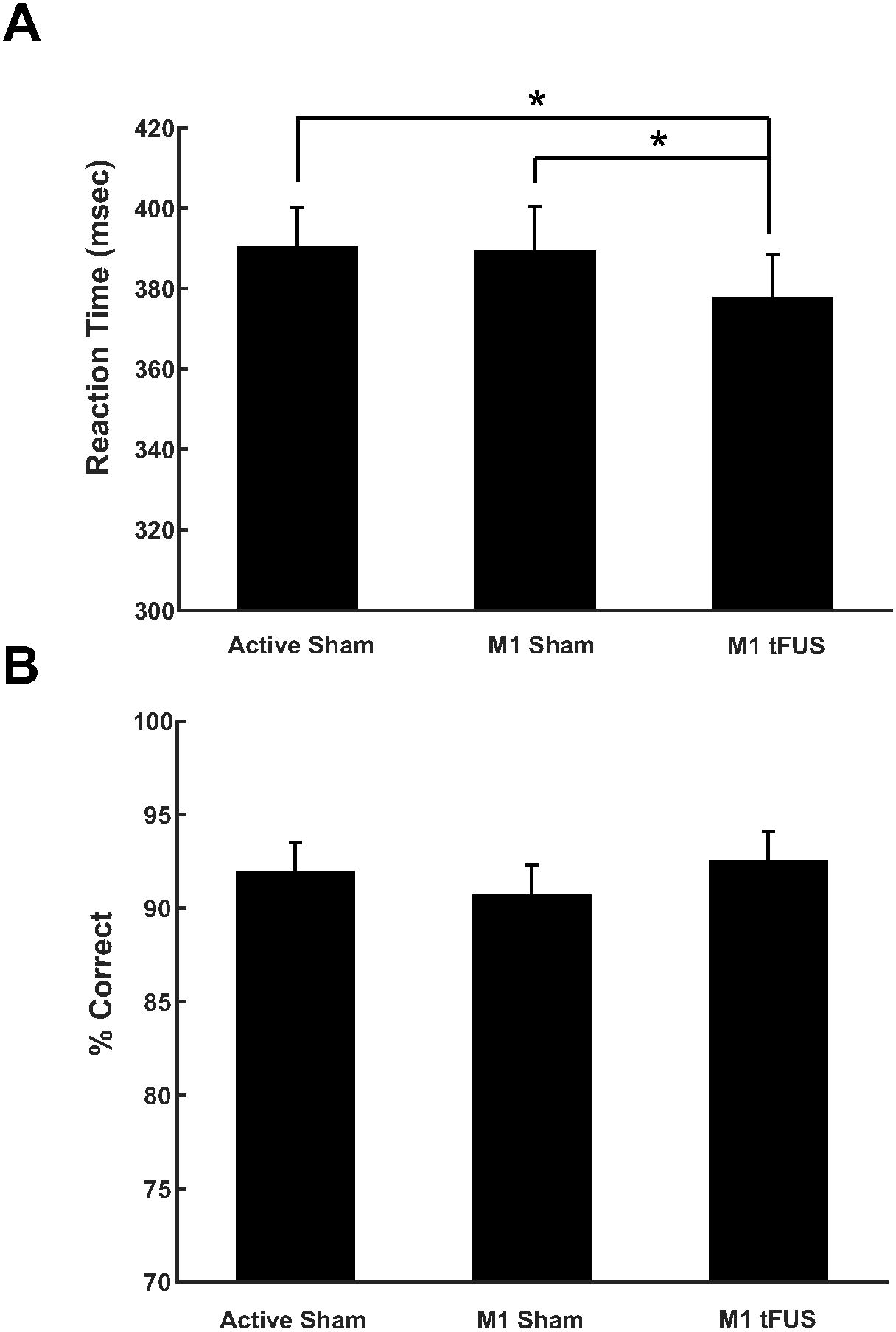
Effect of ultrasound on motor behavior. **A.** Group average (N = 28) reaction time ± SEM in milliseconds (msec) to a stimulus response task. Active sham represents ultrasound delivered to the vertex. M1 sham represents transducer over primary motor cortex (M1) but no ultrasound. M1 tFUS is transcranial focused ultrasound to primary motor cortex. * denotes p < 0.05. **B.** Data from catch trials during the reaction time task. Data is represented as percent correct ± SEM.

## Discussion

In the present study we introduce a novel way to assess the effect of ultrasound on motor cortical excitability in humans by pairing it with TMS, thus allowing for quantification of ultrasound effect on measures of the TMS invoked motor evoked potential. We first demonstrate that concentric and concurrent delivery of TMS and ultrasound (TUMS) is safe in that the energy fields do not interact. Using this pairing, we tested the effect of ultrasound on TMS single pulse recruitment curves, short interval intracortical inhibition and intracortical facilitation as well as on the duration of the cortical silent period on a preliminary small sample. We found ultrasound to attenuate the single pulse recruitment curves. Ultrasound did not further attenuate short-interval intracortical inhibition but significantly attenuated intracortical facilitation. We found no effect of ultrasound on the duration of the cortical silent period. We also tested if the attenuation found for single pulse TMS lasted beyond the period of ultrasound neuromodulation. We found that 500 milliseconds of pulsed ultrasound only attenuated the time point when TMS and ultrasound were delivered concurrently. Finally, we found that ultrasound directed at the primary motor cortex representation of the muscle performing a simple stimulus response task significantly reduced reaction time. This is the first study in humans to assess the effect of ultrasound to the primary motor cortex and indicates that ultrasound largely conveys an inhibitory effect that confers a behavioral advantage.

### TUMS coil

We found no interaction of the ultrasonic sound field with the magnetic field produced by the TMS coil. We also found no initial or persistent effect of concurrent and concentric ultrasound with TMS on either of the hardware after repeated testing. The current TUMS coil design increases the spread and weakens the induced electric field in the head by roughly 30%, though the area of highest current is still located directly beneath the intersection of the TMS coil windings that is concentric with the ultrasound field, making the TUMS design effective for testing spatially specific motor cortical excitability. This reduction in induced electric field in the head necessitates higher TMS stimulator outputs to reach motor threshold for producing a recordable peripheral motor evoked potential. The TMS stimulator output for achieving motor threshold was in the range of 65 – 80% of stimulator output. This is an issue though for participants with naturally high motor thresholds, as this may preclude them from participating in experiments that require TMS at significantly greater stimulator output levels with our TUMS offset than their resting motor threshold level as this may exceed the TMS stimulator output as for example in the cortical silent period study where 120% of motor threshold was used. TMS stimulator outputs of 120 – 130% of resting motor threshold are common for certain TMS protocols and may not always be achievable with the current TUMS configuration. This issue can be minimized with lower profile ultrasound transducers (we used 1.25 cm) or by placing the transducer in plane with the TMS coil windings though TMS wing separation also reduces the induced electric field in the head.

### Mechanisms of action

It is difficult to assess the exact neuronal population that ultrasound affects in humans non-invasively but the use of the TUMS device allows for some inference into the neuronal populations affected by ultrasound as single pulse and paired pulse TMS have been demonstrated to probe different microcircuits in the primary cortex (Di Lazzaro et al. 2012). We used a TMS pulse that produced a posterior to anterior induced current in the brain that has been demonstrated to preferentially target more superficial monosynaptic cortico-cortical neuronal connections in cortical layers two and three that project onto corticospinal neurons producing the so-called I-wave as recorded from spinal electrodes (Di Lazzaro, Oliviero et al. 1998). As such, ultrasound may serve to inhibit these populations resulting in the observed MEP inhibition. However, at high TMS stimulator outputs there is the possibility of initiating additional components of the motor micro-circuitry including direct activation of the pyramidal tract neurons. We found increasing attenuation effects with higher stimulation intensities and thus it is possible for ultrasound to exert its inhibitory effects directly on the pyramidal tract neurons in layer four. TMS has limited penetration depth and its effects are mainly limited to more superficial layers of the cortex, though deeper stimulation is possible with increasing stimulator output at the expense of focality (Deng, Lisanby, Peterchev 2014; Windhoff, Opitz, Thielscher 2013). Focused ultrasound (depending on the orientation of the gyrus to the axial field) can target deep cortical layers as well as sub-cortical targets (Ai, L. Mueller, J.K. Bansal, P. Legon W. 2016; Legon et al. 2014), so it is possible for ultrasound to directly modulate pyramidal tract neurons. It is unclear if the TMS stimulator intensities we used were high enough to produce this direct activation of the pyramidal tract neurons, though we cannot rule out that ultrasound serves to suppress activity directly at the pyramidal cell axon or cell body. The site of mechanism seems likely to be synaptic given the findings of (Kubanek et al. 2016; Prieto et al. 2017) that have shown ultrasound to affect ion channels embedded within cellular membranes and that ultrasound does not appear to have an effect on myelinated axons in vivo (Wright et al. 2017). The fact that we exclusively find inhibition suggests influence of ultrasound on gamma-aminobutyric acid (GABAergic) neurons; however, the specific GABA(A) modulator lorazepam, served to suppress MEP amplitude though only affected motor microcircuitry that produces the late I-waves but not the monosynaptic I wave (Di Lazzaro et al. 2000). Thus, it is unlikely that ultrasound directly affects GABA(A) receptors. The paired-pulse experiment supports our single pulse findings and the contention for ultrasound not to directly affect the circuitry producing I-waves. We employed the method of Kujirai et al. (1993) and did not find a statistically significant attenuation of MEP amplitude for the known ISIs (1-5 msec) to produce short interval intracortical inhibition. (Di Lazzaro, Restuccia et al. 1998) specifically showed for the SICI protocol to suppress the late I-waves. Thus, ultrasound’s mechanisms of inhibition likely do not directly act on GABAergic neurons as does lorazepam for example as it increases SICI (Di Lazzaro et al. 2000). There is evidence for some suppression from ultrasound at SICI inter-stimulus intervals and we posit that perhaps ultrasound may not be an efficient or as strong a mediator of inhibition that could not further suppress the already well-suppressed state of the motor microcircuitry from the SICI protocol that is directly mediated through GABA(A). This could be due to mechano-chemical or mechano-electrical inefficiencies for neural conduction perhaps due to smaller mechanosensitive receptor densities. Despite mostly theoretical proposals of the putative mechanisms of ultrasound (Krasovitski et al. 2011; Plaksin, Kimmel, Shoham 2016; Tyler 2012), the mechanical effects are likely not as robust as direct electro-chemical effects on cellular membranes and/or embedded receptors. There are specific mechanosensitive ion channels in the nervous system (Brohawn 2015; Coste et al. 2010) that have been demonstrated to be sensitive to ultrasonic perturbation (Kubanek et al. 2016; Prieto et al. 2017) though the proliferation and density of these in the human brain is not well understood. One interesting possibility to help explain the mechanism of ultrasonic inhibition is for influence of the ultrasonic pressure field on glial cells (Jordao et al. 2013). Astrocytes express mechanosensitive channels and are known to swell in pathological states. This swelling induces depolarization that may be explained by the opening of chloride channels (Kirischuk 2009). It may be that the negative pressure produced by ultrasound induces astrocyte swelling and chloride release which may explain the general inhibitory effects we see that mimic GABA as the GABA receptor also selectively conducts chloride through its pore. Indeed, one theoretical framework for describing ultrasound effects posits the negative pressure of the ultrasonic wave to expand the lipid membrane (Krasovitski et al. 2011). This may also explain why we didn’t show statistically significant increases in SICI as this is directly mediated by GABA and the contribution of the astrocytes to this would likely be less robust. In the case where there is ample room for inhibition as in the ICF protocol, ultrasound shows robust inhibition and the astrocyte hypothesis may indeed contribute to this effect as lorazepam does not affect intracortical facilitation ((Di Lazzaro et al. 2000), supporting that ultrasound does not directly affect GABA(A) receptors. Unfortunately, the micro-circuitry involved in the ICF protocol is not well understood. ICF does not affect the amplitude or number of descending corticospinal waves despite producing an increase in MEP amplitude suggesting alternate or long-ranging micro-circuitry as compared to single pulse TMS and SICI mechanisms (Di Lazzaro and Ziemann 2013). Ultrasound served to suppress the effect of ICF though not to baseline levels suggesting that, regardless of the specific neuronal population, ultrasound does not serve to totally block excitation but rather its mechanism of inhibition diminishes this effect perhaps by producing a more general state of inhibition through non-localized astrocyte modulation that render the neuronal populations responsible for ICF less receptive to excitation. Additionally, we did not find ultrasound to affect MEP threshold. This is perhaps not too surprising as an increase in TMS output, while increasing evoked potentials, does not affect the threshold for activation. This suggests that the neural populations activated by posterior to anterior TMS induced electric fields have a constant threshold that is unaffected by altered synaptic activity (Di Lazzaro and Ziemann 2013) and ultrasound did not affect this property of the motor micro-circuitry.

### Comparison to motor cortex animal studies

There are numerous reports in mice (King et al. 2013; Mehic et al. 2014; Tufail et al. 2010) as well as in rats (H. Kim, Chiu, Lee, Fischer, Yoo 2014; Younan et al. 2013), rabbit (Yoo et al. 2011) and sheep (Lee et al. 2016) for ultrasound directed at the motor cortex to result in peripheral EMG activity. This de facto represents pyramidal tract neuron excitation. At no point during any of the experiments conducted here did we find ultrasound to spontaneously elicit EMG activity. It is not currently clear why this is possible in animal models and not here, but may be due to the parameters used, the size of the skull relative to the pressure field and/or perhaps due to other indirect considerations (Guo et al. 2017). Specific parameters have been titrated in these small animal preparations that show preference for either excitation or inhibition (H. Kim et al. 2014; King et al. 2013; Yoo et al. 2011). The waveform we employed has previously shown only inhibition (Legon et al. 2014; J. Mueller et al. 2014). It is possible with changes in the duty cycle, amplitude or duration, excitation could be achieved in human motor cortex. Small changes in different parameters (also based on preparation) also may explain the difficulty in producing robust excitation (Wright, Rothwell, Saffari 2015; Younan et al. 2013). Of more importance, may be the effect of skull size or cranial volume relative to the ultrasound pressure field. In recent work, we showed how skull size impacts the intracranial pressure field such that smaller skulls interact with the main acoustic beam more (J. K. Mueller et al. 2017). Previous numerical simulations of experimental setups with rats have found substantial intracranial pressure increases in rat skulls compared to the corresponding transducer’s free-field pressures (Younan et al. 2013). There is greater wave confinement in smaller skulls that will result in stronger and more numerous interactions which could explain motor responses in smaller animals. Indeed, King et al. (2013) demonstrated ultrasound activation to be a function of the product of duration and amplitude and Kubanek et al. (2016) showed the ultrasound effect to be a function of pressure. Thus, it is possible that our waveform combined with our relatively low intracranial modeled pressure of ~ 120 kPa is not sufficient for neural excitation.

We did not find any evidence for ultrasound to affect the duration of the cortical silent period or the amplitude of the MEP during a tonic contraction in our small preliminary (n = 4) sample. Further research should be conducted to definitively determine potential effects. However, a lack of a preliminary effect may be related to alternative motor circuitry underlying the CSP that ultrasound does not affect. There is evidence that the CSP reflects long-lasing GABA(B) mediated inhibition as the administration of baclofen (a specific GABA(B) agonist) resulted in CSP lengthening (Siebner et al. 1998). The circuitry responsible for the silent period is not expressly known but likely has both cortical and spinal components (Wolters, Ziemann, Benecke 2008). Perhaps again, it is the case that ultrasound does not appreciably contribute to an already inhibited state similar to our SICI findings due to less robust mechanisms. We also did not see any effect or a trend for tFUS on MEP amplitude during contraction. Inhibition of MEP amplitude similar to the single pulse or paired pulse data could be hypothesized during contraction as previous research has demonstrated a subthreshold conditioning pulse to reduce MEP amplitude independent of CSP duration (Trompetto et al. 2001) supporting the idea that MEP and CSP are generated by different mechanisms (Wolters et al. 2008).

There looks to be a trend for the inhibition to last beyond the offset of ultrasound though further testing is needed to confirm this. It is currently unclear if the neuromodulatory effects of ultrasound persist beyond the stimulation duration in humans and what the relationship to the length of sonication is to the duration of effect. In Legon et al. (2014) we showed no cumulative effects of ultrasound on the somatosensory evoked potential amplitude though due to the long inter-stimulus interval used (average of 8 seconds) we cannot conclude that the effects of ultrasound neuromodulation persisted to some point out to 8 seconds beyond sonication. As such, we used the TUMS protocol to test this out to 10 seconds beyond offset of ultrasound neuromodulation. We only found statistically significant inhibition of MEP amplitude when the TMS single pulse was delivered during ultrasound though there looks to be a trend for the suppression to extend to 500 milliseconds beyond the offset of ultrasound. We delivered ultrasound for 500 milliseconds and as such, there may be a 1:1 relationship between ultrasound duration and effect duration similar to some repetitive TMS protocols (Fitzgerald, Fountain, Daskalakis 2006). There is evidence in the literature for persistence effects of ultrasound. In rats, (H. Kim et al. 2015) found both their inhibitory and excitatory protocols to last at least 150 seconds post sonication and in one case for an elevated visual evoked potential amplitude to persist at 300 seconds after sonication. However, it is not expressly clear what the duration of sonication was in this study. Additionally, Yoo et al. (2011) also demonstrated persistent effects of their inhibitory protocol where the visual evoked potential amplitude was suppressed for up to ten minutes post sonication after 18 seconds of sonication. Despite the different preparations and parameters, it does look like ultrasound neuromodulation can extend beyond the duration of sonication. The TUMS method is particularly suited to test this as multiple consecutive stimuli are not needed for averaging as in an evoked potential (Yoo et al. 2011; Legon et al. 2014) allowing for precise timing of the sonication relative to the TMS test stimuli in individual trials. We present preliminary work here, though further work should collect more participants and alter durations and parameters to effectively determine the length of ultrasound neuromodulation.

### Reaction Time

We previously found ultrasound with identical parameters as used here to inhibit activity of the primary somatosensory cortex that also resulted in a behavioral improvement in tactile sensitivity (Legon et al. 2014). Here, we also found a behavioral advantage to this ultrasonic neuromodulation concomitant with physiological inhibition. We are confident this behavioral effect is not due to extraneous attentional or potential cueing mechanisms from the ultrasound as we employed an active sham in addition to the passive sham that demonstrated no effect on reaction time. In addition, the data on the catch trials did not show any differences, again demonstrating the effect was not due to attentional factors. It may be paradoxical at first thought that physiological inhibition confers a behavioral advantage but it is a common finding for the balance of excitation and inhibition (or loss of inhibitory tone) to underlie some behaviors (Legon et al. 2016) and neurological conditions such as schizophrenia (Lewis, Hashimoto, Volk 2005). There is evidence in the human motor cortex for inhibition that is necessary for fractionated finger movement that has been demonstrated to be dysfunctional in disease states such as focal dystonia (Hallett 2011). Perhaps ultrasound provides for a mechanism that acts like surround inhibition to sharpen the inputs to the specific finger representation that aids performance. Single finger movements are never completed isolated contractions of one muscle but need simultaneous control of the entire hand and forearm involving the contraction of many muscles acting on different fingers and joints (Beck and Hallett 2011; Schieber 1991). As such, perhaps ultrasound serves to suppress the inputs of the other muscle representations onto the effector, increasing the fidelity of the response signal to this effector and reducing transmission and execution time. The mechanisms of surround inhibition are largely mediated through GABAergic transmission, that from our single and paired pulse experiments ultrasound looks to at least indirectly affect.

## Conclusions

We present here a new method to probe the effect of ultrasound on human motor cortical excitability. The TUMS device allows for a quantifiable method to probe certain micro-circuits within the human motor cortex non-invasively to better understand what circuits and how ultrasound exerts in neuromodulatory effects. We demonstrate a specific ultrasound waveform to induce robust inhibition though the TUMS method can allow for a variety of iterations of ultrasound parameters combined with a variety of TMS protocols to accurately define the circuity ultrasound effects; what changes in parameters do to motor excitability, lending further evidence to elucidate the mechanisms of ultrasound. TUMS is also particularly suited for examining the duration of ultrasound effect on a trial to trial basis and with further work should provide for rich datasets exploring the effect of different parameters. Optimization of ultrasound for specific duration of effects will aid in the ability of ultrasound to be adopted for transient neuromodulation to aid in human brain mapping efforts and eventual application to clinical populations for specific targeting and assessment of brain pathology.

